# Illuminating root-soil mechanics

**DOI:** 10.1101/2024.10.02.616056

**Authors:** James Le Houx, Daniel McKay Fletcher, Alberto Leonardi, Katherine A. Williams, Nancy Walker, Fernando Alvarez-Borges, Ebrahim Afsar Dizaj, Madhu Murthy, Ronan Smith, Liam Perera, Navid Aslani, Andrew James, Sharif Ahmed, Tiina Roose, Siul Ruiz

**Affiliations:** ISIS Neutron & Muon Source, Rutherford Appleton Laboratory, Didcot, OX11 0QX, United Kingdom; The Faraday Institution, Harwell Science and Innovation Campus, Didcot, OX11 0RA, United Kingdom; University of Southampton, Faculty of Engineering and Physical Sciences, University Road, Southampton, SO17 1BJ, United Kingdom; Scottish Rural University College, Rural Economy, Environment and Society, West Mains Road, Edinburgh, EH9 3JG, United Kingdom; Diamond Light Source, Rutherford Appleton Laboratory, Didcot, OX11 0QX, United Kingdom; University of Portsmouth, Faculty of Science and Health, King Henry I Street, Portsmouth PO1 2DY, United Kingdom; Yale University, School of the Environment, New Haven, CT 06511, USA; University of Adelaide, Faculty of Health and Medical Sciences, Adelaide, SA 5005, Australia

**Keywords:** Multimodal X-ray computed tomography, X-ray diffraction mapping, plant root growth, soil bioturbation, finite element modelling

## Abstract

Soil compaction and escalating global drought increase soil strength and stiffness. It remains unclear which plant root biomechanical mechanisms/traits enable growth in these harsh conditions. Here, we combine synchrotron X-ray computed tomography with spatially resolved X-ray diffraction to characterize the biomechanics of a replica root-soil system. We map the strain field around the root tip, finding strong agreement with finite element simulations, thereby demonstrating a promising new in-vivo measurement protocol.

## Main

Rising frequency and severity of droughts threaten global food security and infrastructure development in much of the world^1^. Soil compaction resulting from intensified farming adversely impacts ∼25% of Europe’s mechanized arable land area^2^, while drought impacts over 3B people globally^3^. Drought and compaction increase the strength and stiffness of soils^4^ creating environments that are challenging for plant roots and burrowing organisms to penetrate. These conditions make it difficult for them to forage for water and nutrients^4^. Plant roots, along with other tunneling organisms, help sustain soil function through soil bioturbation^4^. Formation of bio-pore networks enhances deep drainage of water mitigating flooding, enabling diffusion of oxygen into deeper soil horizons, and priming soil microbial activity^4^. These effects create favourable conditions for agricultural systems and can mitigate desertification.

Plant roots can penetrate surprisingly rigid substrates, but there are limits to this ability. The localized mechanisms behind this process are not well understood, as roots penetrate soil through a complex ensemble of multi-scale processes. Root turgor pressure allows the root to axially extend into the soil, while cells near the root tip multiply and reorient themselves through an enzymatic process that reduces local friction^5^ (Figure 1 A). Although these processes allow roots to exert enough pressure to fracture solid chalk^6^, there is no clear understanding as to how or where these pressures are applied. Root caps have been observed swelling in compacted soils, which provides mechanical advantages for the subsequent growth^7^. However, it is unclear whether this is because of loosening the forefront of soil or extending pre-existing faults in the media (e.g. cracks).

**Figure 1.**
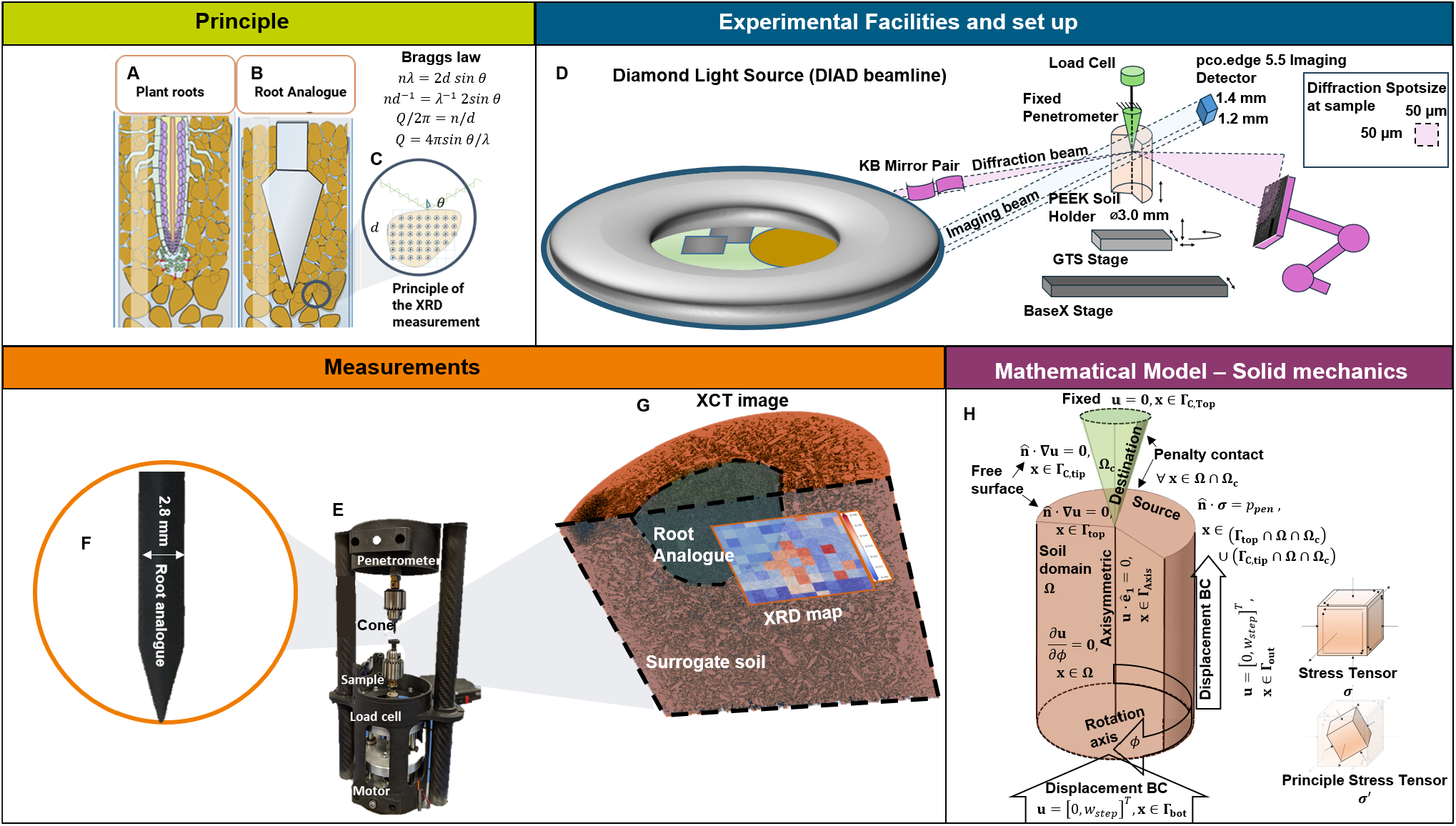
Methods, experimental set up and mathematical model. The method is being developed for A – measuring biomechanics associated with root growth by using B – cone penetrometer root analogues for C – correlated XCT and XRD. D - Diamond Light Source and diagram of the experimental setup for the correlated XCT and XRD measurements at the DIAD beamline. E – Developed in-situ compression rig with F - the cone penetrometer and sample near the detector using a root analogue cone. G - Overlaid XCT and XRD result from DIAD - XCT 3D image and 2D diffraction map representing the ensemble of inferred stress values across the sample. H - Diagram of the mathematical mechanical model domain and the associated boundary conditions simulating the experiment.

Recently, researchers have used X-ray computed tomography (XCT) to visualize soil displacements associated with growing plant roots on the order of 3-10 µm resolution under loosely packed soil conditions^8^. However, soil deformation itself is more difficult to quantify under more compact soil conditions, which is expected to become more prominent in agricultural settings given land use intensification and increased machinery weight^2^. Most conventional laboratory tests (e.g. triaxial) can only deliver insight on dense soil deformation under simplified stress conditions and at the macro/bulk scale. They cannot directly observe stress states in localized areas of deformation, such as shear bands^9^.

X-Ray diffraction (XRD) has emerged as a promising tool for inferring the mechanical deformation state of materials^10^. This non-destructive technique uses the structure properties of a material’s crystals as a strain gauge. Under applied loads, variations in a material’s structure lattice parameters can be determined based on Bragg’s Law^11^(Figure 1 C). Given a material with known elastic constants, local strains can be used to infer corresponding stresses. This method has shown success in quantifying stress fields in granular media via individual particles under confined uniaxial loads^12^, but has not to date been applied to understanding localized biomechanical processes in soil.

This work shows proof of principle for applying coupled multimodal XCT and XRD measurements to quantify the strain and infer the mechanical stress associated with root growth in soil. We used an artificial root model (i.e., cone penetrometer Figure 1 B) moving into a surrogate soil material, gypsum (calcium sulfate dihydrate, *CaSO*_4_ *·* 2*H*_2_*O*). All measurements were conducted at Diamond Light Source using the K11 Dual Imaging and Diffraction (DIAD) beamline^13^. A custom designed load frame was engineered to simulate root growth into the soil (Figure 1 E), using a conical tip (Figure 1 F) as the loading tool. The load frame used a stepper motor to drive the sample upward into the conical tip, penetrating the sample whilst a load cell measured the forces exerted during penetration. Gypsum was selected as the surrogate soil for its relatively soft crystalline structure, facilitating the diffraction measurements. We prepared the sample holders with a wet paste of soil and allowed the samples to dry, thereby emulating strong capillary potentials that are representative of drought conditions. Once the sample was loaded, XCT and XRD measurements were taken sequentially (Figure 1 G), where XRD probed the strain field by measuring the gypsum peak shift as a function of the diffraction probe position.

The cone penetrated progressively into the sample to four different depths – 0 mm, 1 mm, 5 mm, and 8 mm. At each depth, XCT measurements visualized (Figure 2 A-D) the evolution of the pore space (Figure 2 E-H). These images highlight the “compression zone”, characterized by a reduction in both the number and size of pores surrounding the cone tip as it penetrated the soil.

**Figure 2.**
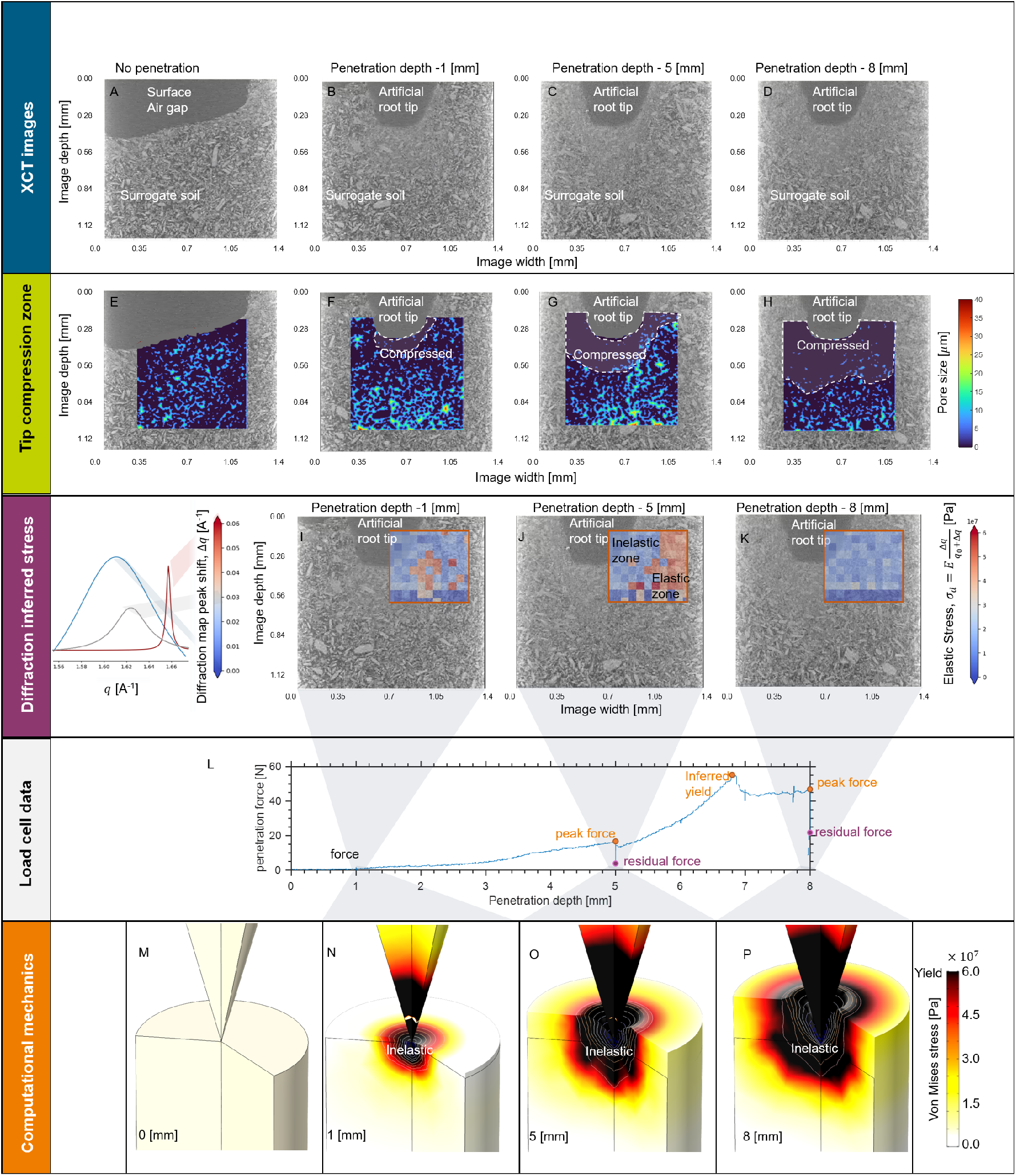
Results of the coupled XCT imaging and XRD during root analogue penetration. A - The surface of the sample without the cone. B - The root analogue conical tip penetrating 1 mm, C - 5 mm, D - 8 mm into the surface. Compression zone quantified by considering the changes in the pore size around the root tip for E - surface, F - 1 mm penetration depth, G - 5 mm penetration depth, and H - 8 mm penetration depth. XRD inferred elastic stresses overlay the XCT slices for I - 1 mm, J - 5 mm, and K - 8 mm depths, where the stresses are inferred from the diffraction peak shifts (040) and the elastic modulus of gypsum. L - Illustrates load cell forces associated with penetration tests. Computational models are used to assess the yielding behavior, where the artificial root penetrates into gypsum continuum from M - the surface, N - 1 mm, O - 5 mm, and P - 8 mm depth, highlighting the expanding yield zone, creating a region that cannot be readily measured via XRD.

While the XCT results support our initial hypotheses, the overlaid diffraction maps revealed an unexpected finding: stress appears to reduce at the highest penetration depth. (Figure 2 I-K) illustrates the diffraction peak (040) shifts associated with each loading step. For the 1 mm load step (Figure 2 I), we monitored a small region just distal to the cone tip, where peak shifts indicative of loading were observed. In contrast, at 5 mm penetration, Figure 2 J reveals a region adjacent to the cone tip with no discernible peak shifts. Beyond this region (∼200 µm), we identify a topographical boundary of diffraction peak (040) shifts. Whereas, at a penetration depth of 8 mm (Figure 2 K), these peak shifts are no longer undetectable within the diffraction window. Notably, the penetration forces measured by the load cell (Figure 2 L) continue to increase up to approximately 7 mm penetration depth, after which progressive material failure results in a stabilized force reading.

To gain fundamental insights into the measurements, we conducted a finite element mechanics digital twin simulation (Figure 2 M-P). The simulation considered a cylinder of gypsum treating it as an elasto-plastic material^14^. The simulation results highlight the evolution of the inelastic region around the penetrating cone tip, where at 1 mm penetration (Figure 2 N) the inelastic zone is very local to the cone tip, but at 8 mm propagates to the outer radius of the cylinder (Figure 2 P). This result provides insight as to why we see an un-shifted zone just distal to the cone tip in the diffraction maps (Figure 2 I-K). When soils deform plastically, soil particles re-orient and/or break forming new particulate arrangements, with mean intra-particle stresses reducing sharply with decreasing particle sizes^9^. Our XRD peak shift measurements reveal that the elastic deformation of gypsum crystal structures indicates that inter-particle interactions (e.g. breakage, reorientation) result in measurable changes in intra-particle strain, suggesting a relaxation of intra-particle elastic stress. This behaviour correlates well with the observed reduction in mean pore size around the cone tip. Therefore, the topographical boundary of XRD peak shift measurements is an indicator of the extent of the plastic deformation zone surrounding the cone.

This method offers a robust approach in understanding fundamental biomechanics associated with root growth, effectively addressing common limitations encountered by researchers studying local mechanics behaviour. Importantly, root tip growth is unlikely to drag surface material to lower depths; therefore, in-vivo diffraction maps are expected to closely resemble those in Figure 2 I-J rather than Figure 2 K.

Controlled experiments with natural plant roots will yield valuable insights into the location of root loads (i.e. radial or axial) and likely reveal whether plant roots are exploiting imperfections in their growth medium (i.e. crack propagation). Imaging data similar to that shown in Figure 2 A-H, will elucidate the morphological features of plant roots that facilitate penetration through compacted soils, whilst simultaneously identifying pre-existing faults/cracks within the soil structure. Furthermore, diffraction maps (Figure 2 L-O) will demonstrate how these growth patterns confer mechanical advantages. Traits that enhance growth through compacted soils can be strategically integrated into cover crop breeding programs for regenerative agricultural practices, informing rewilding strategies for over-farmed/desertified areas and ultimately contributing to ecosystem resilience. Beyond sustainable agriculture and land management, our results have broader applicability in material science and geotechnical engineering, particularly in addressing the complexities of characterizing plasticity in soil-structure interactions. Here, we demonstrate that correlated multimodal XCT and spatially resolved diffraction measurements can effectively characterize these interactions, aiding in the precision design of foundations for critical infrastructure, such as pile-supported wind turbines^15^.

## Methods

### Dual Imaging And Diffraction (DIAD) beamline

The Dual Imaging and Diffraction (DIAD) beamline at Diamond Light Source is a new instrument designed for quasi-simultaneous space-time correlated imaging and diffraction experiments. DIAD combines quasi-simultaneous (sequentially up to a frequency of 10 Hz) X-ray Computed Tomography (XCT) and X-ray Diffraction (XRD) techniques using a dual-beam setup. Both beams are spatially registered, enabling ‘image-guided diffraction’, and allowing for the acquisition of both morphological and structural/compositional information from samples in real-time. XCT offers high-resolution 3D images of the macro-soil structure, revealing the evolution of the root systems and soil morphology. Concurrently, XRD provides information on the crystalline phases, composition and strain within the soil and root materials. This dual approach allows for the correlation of physical structure with material properties for dynamic systems, improving understanding of the mechanical and compositional changes occurring during root growth.

DIAD uses a novel optical layout where the insertion device, a fixed-gap hybrid wiggler, generates a wide photon fan that is split into two independent beams, each passed through an independent series of optical elements and subsequently combined at the sample position. The imaging beam is full field (1.7 mm × 1.7 mm) and can be configured in pink beam mode, where the spectral profile is determined by a series of mirror stripes and filters, or monochromatic beam, where a pair of cryo-cooled Si (111) double-bounce crystals (DCM) are used to monochromate the beam in the energy range of 7-38 keV. For this study, the imaging branch was set in pink low-angle beam with a Pt-Pt mirror stripe setup and a 4 mm Al filter. The diffraction beam operates in monochromatic mode, also using a DCM, but has an additional optical element of a Kirkpatrick-Baez (KB) mirror pair. The KB mirror system focuses the diffraction beam to a spotsize of 50 µm × 50 µm at the sample position, and which can then be scanned across the imaging Field Of View (FOV).

The imaging detector is a pco.edge 5.5 camera with a 10 µm thick terbium-doped Gadolinium Gallium Garnet (GGG:Tb) scintillator, featuring a pixel-size of 0.54 µm and an imaging FOV of 1.4 mm *×*1.2 mm. The diffraction detector is a cadmium telluride Pilatus2 × 2M large 2D area detector, mounted on an industrial robot arm. The imaging detector is mounted downstream of the incident photon beam, whilst the diffraction detector is positioned off the main axis of the imaging beam path, capturing partial Debeye-Scherrer rings. This configuration allows both detectors to operate simultaneously without needing repositioning or causing interference. In order to correct for the changing beam-to-sample geometry of the moving diffraction beam, a geometry calibration scan is performed using a *LaB*_6_ standard, mapped over the imaging FOV on a 2D grid. The diffraction geometry is then automatically corrected for in the azimuthal integration step of the data reduction process in the open-source software, DAWN Science.

### In-situ compression rig emulating root penetration

The process of root penetration into soil during growth is analogized by a cone penetrometer. This set-up effectively captures the displacement of soil material as it accommodates the penetrating object, whilst the resistance encountered by the cone provides insights into the mechanical properties. Unlike natural root growth, the cone penetrometer allows for controlled displacement and force profiles, establishing a reliable stress-strain field for characterization. Additionally, root penetration is a biological process influenced by factors such as root growth rate, soil moisture, and nutrient availability, whereas the cone penetrometer operates mechanically. Consequently, the penetrometer can follow a pre-determined displacement to reduce the effects of spatial aberration of the photon beam, whereas roots alter their growth direction in response to soil conditions.

XRD is employed to characterize the strain within surrogate soil materials. Surrogate soil was chosen over natural soils due to their compositional and structural/microstructural homogeneity, as well as facilitating diffraction measurements through the crystalline structure. This homogeneity facilitates the mechanical interpretation of peak shifts and broadening during diffraction measurements. Despite these differences, surrogate soils retain a similar morphology to natural soil. Analysis of the diffraction patterns gives access to strain distribution information within the soil samples. This involves collecting ensemble or averaged data to obtain a comprehensive view of the strain characteristics across the soil system in a 2D map. The diffraction data provides insights into the mechanical stresses experienced by the soil during root penetration or cone penetration experiments.

The cone penetrometer experiments utilized three moving sample stages. The baseX stage was responsible for positioning the specimen within the FOV. The general tomography stage (GTS) was used for alignment and rotation. The penetrometer stage, controlled by a linear actuator, loaded the surrogate soil against the fixed penetrometer, ensuring consistent imaging FOVs. The compression rig featured a Nanotec captive linear actuator and a high-resolution magnetic incremental encoder to move the vertical stage, along with an HBM U9C full bridge 100 N strain gauge serving as the load cell. The frame is constructed with carbon fibre support pillars and steel mounting plates, allowing the X-ray beam to pass only through the carbon components to minimize photon attenuation. Additionally, two articulating chucks were used to align the root analogue and sample holder and subsequently locked in position. The compression rig was directly integrated into the beamline control layer to automate the experiment.

A polyether ether ketone (PEEK)-designed cone with an outer diameter of 2.8 mm was mounted to a load cell fixed at the bottom of the frame. We note that we conducted tests with the load cell fixed at both bottom and top, and the results were in agreement. PEEK was chosen for its relative X-ray transparency but sufficient mechanical properties. For our experiments, we selected gypsum (calcium sulfate dihydrate, *CaSO*_4_ *·* 2*H*_2_*O*) as the surrogate soil material. In the sample preparation lab, soil samples were created by filling a small PEEK sample holder (14 mm depth, 3 mm diameter) with soil. Dried paste samples were loaded to emulate soil conditions under drought. 4 g of gypsum powder were mixed with 2 g of water to form a paste, which was then slowly injected into the sample holder using a syringe, filling from the bottom to the top. A diagram of the experimental setup is given in Figure 1 D-E.

### Automated acquisition and processing of correlated measurements

The experiments were conducted using both XCT and XRD sequentially. Full tomograms were captured using a pink beam with a peak flux of 31 keV and an energy band width of approximately 15 keV. Tomograms were acquired with the root at the centre axis of the FOV, and root tip 750 pixels from the top of the FOV. The exposure time was 0.014 s, with 5000 projections over a 180° range. Additionally, 50 dark and flat field images were taken for dark/flat field correction. The image resolution was 2560 pixels × 2560 pixels × 2160 pixels. XRD data were collected at an energy of 22 keV and a focused spot-size of 50 µm × 50 µm across a sub-window of the FOV (coordinates: 180 pixels to 2280 pixels depth, 1280 pixels to 2280 pixels width) around the cone. Diffraction maps were generated at coarse (10 × 11 grid) and fine (20 × 22 grid) resolutions, with each point measured for 4 seconds. Coarse maps took approximately 7 minutes, while fine maps took just under 30 minutes. XCT and XRD measurements were taken at four penetration depths, provided the load cell force remained below 60 N, chosen to reduce the likelihood of load cell calibration drift. Simultaneous force measurements were recorded during the whole experiment using the integrated load cell.

The measurements were obtained using an automated acquisition and processing pipeline. The linear actuator was programmed to move to the specified penetration depth, initiating sequential tomography and diffraction mapping. During tomography acquisition, an automated reconstruction pipeline was triggered using the open-source Python software, Savu ^16^, with complete reconstruction pipelines available in the online Pure repository. For diffraction, an automated data reduction pipeline in Dawn Science ^17^ was employed to convert each 2D detector image into a 1D diffraction pattern, correcting for variable sample-detector geometry using a *LaB*_6_ calibration at the start of the experiment; this reduction pipeline is also documented in the Pure repository. After completing the tomography and diffraction maps, the linear actuator advanced to the next penetration depth, repeating the process until the entire sample dataset was acquired.

### Post-processing of XRD data

The azimuthal integrated XRD data were used to infer 2D elastic strains based on changes in the peak position in the *q*-space. A 10 by 11 grid overlaying the projected area/radiograph of the sample (root tip and soil) was used to take XRD measurements at each position of the grid, representing an ensemble of diffraction measurements through the sample volume (Figure 1 G). We monitored changes of peaks from an initial *q* value of *q*_0_ = 1.61 Å^*™*1^ as determined from an unloaded sample. A gaussian distribution was fit to the and the location of the maximum amplitude value was used as the peak *q* location. For the subsequent loading steps where the relative sample position where the root had penetrated 1 mm, 5 mm, and 8 mm into the soil depths, XRD maps were generated. Differences were taken for each step and the initial diffraction map, and the strain was estimated by

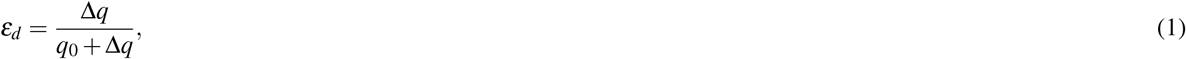

where Δ*q* is the is the change in the peak location in the penetrated samples (i.e. 1 mm, 5 mm, and 8 mm) with respect to the initial state of the sample (0 mm). We considered the soil sample to have material properties associated with a dried gypsum paste, where the elastic modulus is *E*_*g*_ = 1.75 MPa ^18^. As such, we are able to infer elastic stress values based on *σ*_*d*_ = *E*_*g*_*ε*_*d*_. We note that the natural gypsum rock has a failure stress at *σ*_*y*_ = 60 MPa ^19^, which was consistent with the resulting XRD inferred stress values we monitored. We note that these stresses are higher than the values obtained from the load cell measurements, as loads close to the cone tip likely induce more concentrated stresses that cause the gypsum to fail locally. Bulk scale imperfections may cause a reduction in the load cell measurements. Scripts for determining peak positions, generating diffraction maps, and taking the differences between maps were all developed in python and are available in the data repository.

### Post-processing of XCT data

For the same data set, XCT images were used to monitor the evolution of the soil pore space over the different penetration steps. Representative 2D central slices were extracted from the XCT data for the 0 mm, 1 mm, 5 mm, and 8 mm penetration steps. Each of these four images were converted into an 8-bit image for subsequent processing. A median filter with a square kernel and radius of 4 pixels was applied to the images to mitigate shot noise and smoothen/average the imaged data. To segment the pore space from the solid soil fraction, we employed Otsu’s thresholding method, which identifies pixel values that maximize inter-class variance based on the image histogram ^20^. Following segmentation, a distance transformation was applied to quantify and label pore sizes. Given the pore space, denoted as Ω_*p*_ (i.e. pixels with a value 0 in the segmented image), and the boundary between pores and solids, *∂* Ω_*p*_ (i.e. pixels with adjacent pixels having 0 on one side and 1 on the other), we determine the smallest kernel that can fit at any pixel location in the pore space 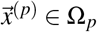 as having a radius defined as:

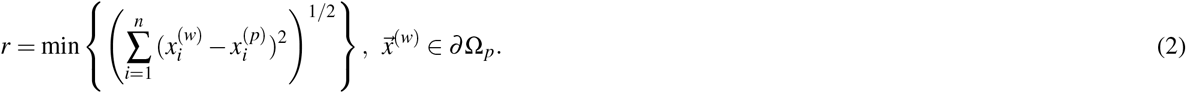

Each pixel in the pore space is color coded according to its radius value. This processing allows us to monitor the evolution of the pore space near the tip of the root throughout the penetration process, facilitating the assessment of bulk scale compression zones. Image processing was carried out in Matlab.

### Modelling elasto-plasticity associated with penetration

To validate XRD results with the established theory of deformation, we modelled the mechanics of cone penetration through a uniform medium. While there is granularity present in the XCT images, the medium is assumed to behave as elasto-plastic bulk material for the simplicity of the model, where we consider an equilibrium equation as:

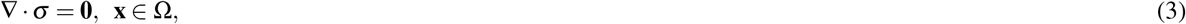

where *σ* = *σ*_*i j*_ represents the stress tensor and Ω is the material in the cylindrical volume analogous to the experiment. For the linear elastic case, we relate the stresses to the strains as

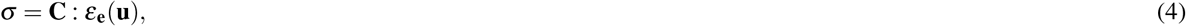

where **C** is the elasticity tensor (4th order) and *ε*(**u**) is the elastic strain tensor (2nd order) defined by:

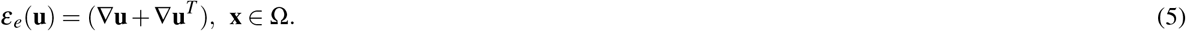

Note that it’s convenient to consider the vector-matrix convention for the constitutive laws (i.e. reshaping the tensors). If we consider *σ* consisting of *n* by *m* stress components, or a 3 by 3 matrix:

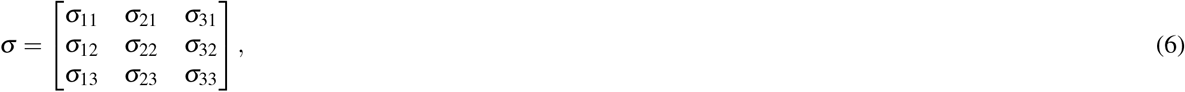

we can collapse them onto a 9 by 1 vector:

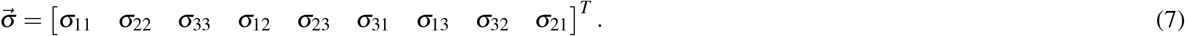

As such, *σ* = **C** : *ε*_**e**_(**u**) can be equivalently expressed as:

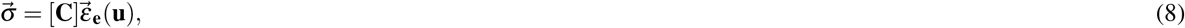

where [**C**] is the matrix representation of the stiffness tensor **C**. In this work, we aim to improve understanding of both the elastic and inelastic deformations, therefore, we break the components of strain into elastic and plastic ^14^

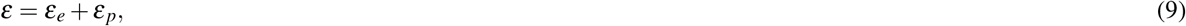

where the elastic strain *ε*_*e*_ is described by eq 5. To quantify plastic deformation, we must assess the material’s yielding, which is best handled by examining the principle stress components. If we consider the stress tensor as a matrix with rows and columns (indexed by *i* for rows and *j* for column), we solve the diagonalization problem:

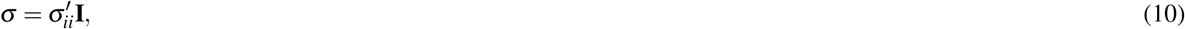

where **I** is the identity matrix and 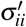 are the magnitudes of the principle stresses in an associated direction, which are defined as the eigenvalues of the stress tensor. Solving 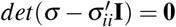 for the different values of 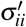 provides the principle stresses. In particular, for 3D Cartesian coordinates, we consider our collection of principle stresses to be

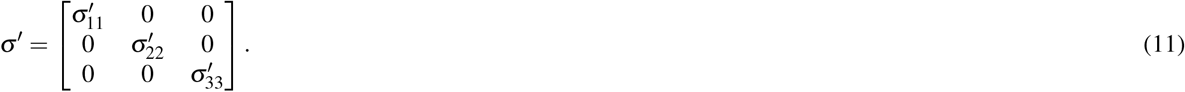

We use the principle stresses to formulate our constituent laws, where we consider elastic deformation for small stresses and perfectly plastic deformation with an associated flow law for stresses which exceed the von Mises yield criterion, defined as:

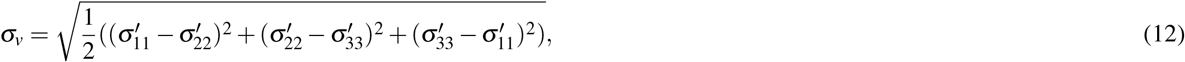

representing a measure of the magnitude of the differences between the principle stresses and indicating the shear stress of the material. We note that in standard engineering practices, Mohr-Coulomb criteria is often invoked to quantify inelastic behavior. However, as we’re dealing with smaller scales and trying to model a dried paste of gypsum, the von Mises yield criteria is sufficient for describing the inelastic behaviour of our system. We define a yield function based on our criteria as:

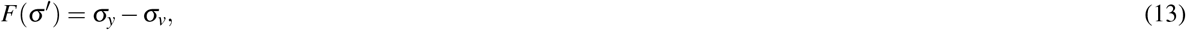

where *σ*_*y*_ is the yield stress of the material. The material will deform in an inelastic manner when *F*(*σ*^*′*^) = 0 which can be interpreted to mean that the stress conditions have a limitation based on the materials ‘strength’. Once that threshold is reached, the material starts to slip, reorient, and effectively flow. The amalgamation of this behavior is categorized as inelastic behavior, which is characterized in this model as plasticity. We note that these processes occur between a materials yield strength and ultimate (i.e. failure) strength. If we focus again on the concept of material ‘flowing’, we should think of a fluid and come to the idea that strain rates are proportional to stresses. Note that the material behaves as incompressible during yield^14^. Note that the flow of material is local within the yield zone. The remote elastic zone is still compressible and acts to constrain the solution. We can, therefore, re-write our constitutive elastic law in terms principle stresses and of rates:

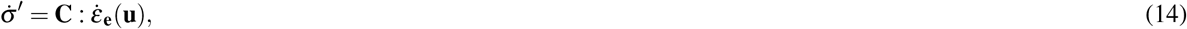

where 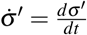 and 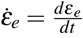 are the stress and elastic strain rate tensors respectively. We revisit the yield function and create a consistency condition at the yield surface (i.e. at *F*(*σ*^*′*^) = 0):

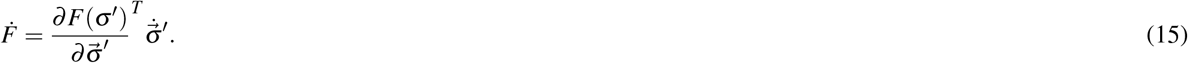

We apply an associated flow rule to quantify the plastic strain behavior, which specifies that the incremental change in the plastic strains are proportional to the changes in our yield function with respect to changes in our principle stresses, expressed as

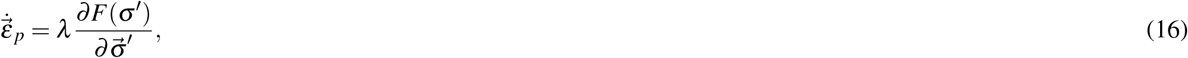

where *λ* is the plastic multiplier, effectively acting as a Lagrangian multiplier that ensures the yield function remains at zero. We consider perfectly plastic conditions for this work, which allows for certain estimates for the plastic multiplier. If we re-write eq. 9 in terms of the elastic strain i.e. (*ε*_*e*_ = *ε ™ ε* _*p*_), substitute this term in for eq. 14 (i.e. 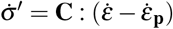)), and substitute in our consistency condition for the eq. 15, we come to:

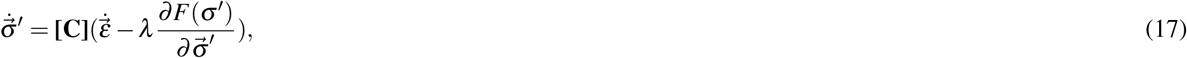

where we note here that we have strain rates as a function of both the stress rates and the stress. We can substitute eq. 17 into the consistency condition eq. 15:

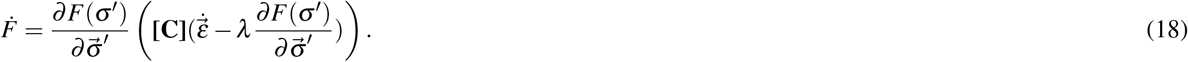

At 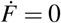, we obtain an expression for the plastic multiplier:

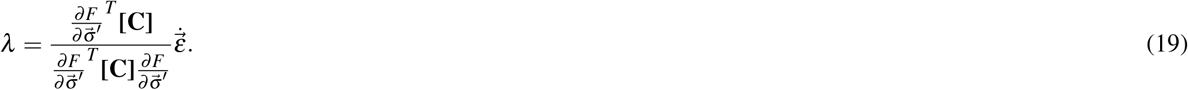

We can substitute this (eq. 19) into eq. 17 and find a common factor for 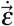:

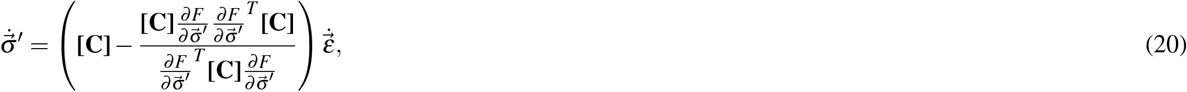

Consequently, we only have stress rates and strain rates (i.e. no absolute stress values). This gives rise to a new effective elasto-plastic stiffness tensor:

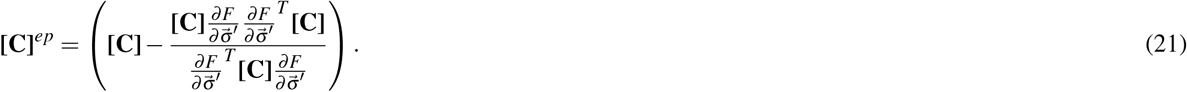

Thus, our domain equations are as follows:

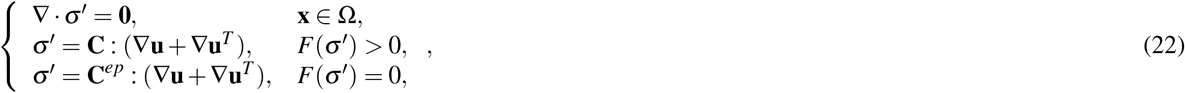

While *λ* can be analytically determined in the elasto-perfectly plastic scenario, this was resolved numerically using COMSOL considering the Kuhn-Tucker conditions.

The work considers a set of boundary conditions. We consider axial symmetry along the central axis and a cylindrical tube of material rising towards a fixed stiff cone (Figure 1 H). Axial symmetry is designated as:

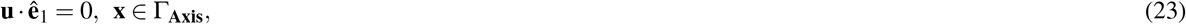

which restricts radial displacements at the origin. Considering an angular coordinate *φ*, we know that there are no changes to the state variable (displacement vector) at any orientation:

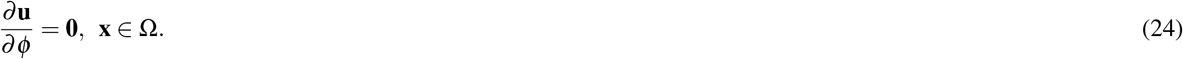

Simulations are carried out by rising the soil (gypsum) domain into the artificial root (cone). The top of the cone is considered fixed:

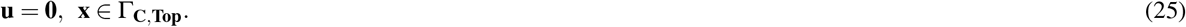

The bottom and edges of the soil domain boundaries have a prescribed vertical displacement:

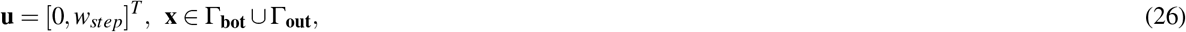

where *w*_*step*_ represents the parametric vertical steps taken during the simulation. For conditions where the surface of the cone tip and the soil are not in contact, these surfaces are considered free surfaces:

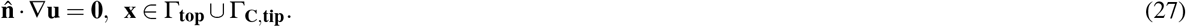

We define contact pairs between the soil and the cone. As the soil domain moves into the cone, we designate the soil surface as the ‘source’ boundary and the cone surface as the ‘destination’ boundary. When the cone domain Ω_**c**_ overlaps with the soil domain Ω (i.e. *∀***x** *∈* Ω *∩* Ω_**c**_, a penalty pressure is applied to the surfaces that are intended to push against one another:

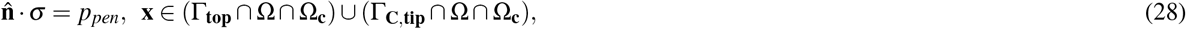

indicating that a penalty pressure is exerted on the surfaces, equal and opposite, in order to simulate the contact. Hence, our final system of equations is as follows:

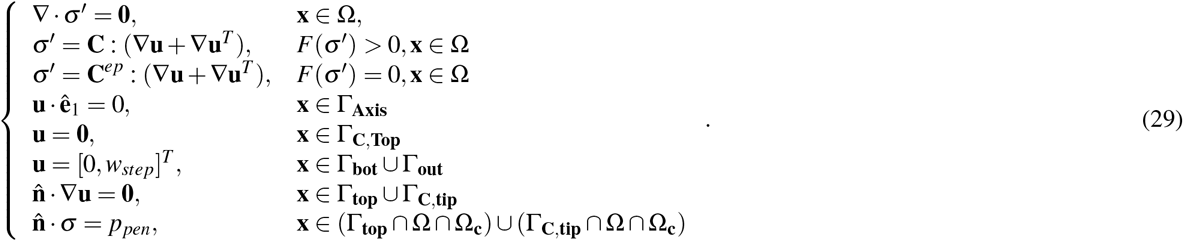

## Data Records

All X-ray diffraction and X-ray imaging data used in this study can be found in the Pure repository: https://doi.org/10.5258/SOTON/D3309.

## Code availability

All scripts used to process the data can be found in the Pure repository: https://doi.org/10.5258/SOTON/D3309.

## Acknowledgements

This work was partially funded by SR’s BBSRC Discovery Fellowship BB/X010147/1 and Royal Society University Research Fellowship URF131622 for “Quantifying soil biomechanics with X-Ray diffraction-imaging and mathematical models”.

This work was partially funded by the Rutherford Appleton Laboratory and Faraday Institution, through JLH’s emerging leader fellowship FIELF001. This work was part funded by the National Research Facility for Lab X-ray CT (NXCT), EP/T02593X/1 and the EPSRC prosperity partnership with Imperial College, INFUSE, Interface with the Future - Underpinning Science to Support the Energy transition EP/V038044/1. DMF: This work was supported by the Rural and Environment Science and Analytical Services Division (SRUC-C5-1). This work was also part funded by TR’s ERC Consolidator grant 646809 (Data Intensive Modelling of the Rhizosphere Processes), BBSRC SARIC BB/P004180/1, BBSRC SARISA BB/L025620/1 and EPSRC EP/M020355/1, which partially funded KW. NW was funded via NERC Inspire DTP awarded to TR. EAD PhD studies are funded via Royal Society Wolfson Fellowship award to TR.

The authors would like to acknowledge the Diamond Light Source for access to the Dual Imaging And Diffraction (DIAD) K11 beamline under proposals MG30961, MG32138 and MG33343. We also extend our sincere thanks to Peter Garland in the preparation and machining of sample holders and to Armando Pueyos for his expertise in the integration and control of the rig.

In memory of Max Leighton (1993-2024), who left this world a better place.

## Author contributions statement

SR and JLH conceived the study. JLH, DMF, and SR designed the experiment(s). JLH, AL, LP, NA, AJ, and SA designed and developed the beam line and load frame. JLH, DMF, AL, KW, NW, EAD, MM, RS, TR, and SR conducted the experiment(s). DMF, AL, KW, and SR analysed the results. DMF, FA and SR developed the conceptual model. All authors reviewed the manuscript.

